# Effect of diapause on cold-resistance in different life-stages of an aphid parasitoid wasp

**DOI:** 10.1101/489427

**Authors:** K. Tougeron, L. Blanchet, J. van Baaren, C. Le Lann, J. Brodeur

## Abstract

To overwinter, insects from mild temperate areas can either enter diapause or remain active. Both strategies involve costs and benefits depending on the environment. In the first case, the emerging individuals will resist winter but have a reduced fitness because diapause entails physiological and ecological costs. In the second case, individuals need to be cold-resistant enough to withstand winter temperatures during their immature and adult stages, but could avoid diapause-associated costs. In mild temperate areas, the cost-benefit balance between the diapause and the non-diapause strategy would likely change in response to climate warming. A trade-off between these two strategies should lead to reduction of diapause expression in some populations. We explored the importance of such trade-off through the comparison of cold resistance capacities among different life stages in diapause and non-diapause individuals in a population of the aphid parasitoid *Aphidius ervi* (Hymenoptera: Braconidae) originating from western France where a decrease of diapause incidence was recently observed. As a proxy measure of insect physiological cold resistance, the Super Cooling Point (SCP) was determined for non-diapausing and diapausing prepupae and adults that went through prepupal diapause or not. Diapausing and non-diapausing prepupae were equally cold-resistant (−24.20 ±0.30°C vs. −24.74 ±0.36°C, respectively), and overall more resistant than adults. Adults that went through diapause as prepupae were less cold resistant (−17.85 ±1.10°C) than adults that have not undergone through diapause (−21.10 ±0.54°C). We also found that diapausing prepupae and adults that have undergone diapause were lighter than their non-diapausing counterparts, at a comparable size, suggesting higher energetic expenses during diapause. These results suggest a trade-off between diapause expression in prepupae and cold resistance and life-history-traits in adults. We conclude that selection could favor insects that do not enter diapause and thus avoid its associated costs while taking advantage at exploiting the mild winter environment.

## 1. Introduction

In temperate areas, insects have evolved various strategies to spatially or temporally cope with recurrent unfavorable periods (Danks, 2006). They can migrate, resist freezing conditions or, as for most insects, avoid lethal freezing of body fluids (Bale, 1987). These freeze-intolerant insects have the capacity to lower the temperature at which their cell fluids crystalize (i.e. the Super Cooling Point (SCP)) by producing cryoprotectant molecules (e.g. polyols, sugars), eliminating ice nucleators in gut contents and partially dehydrating their tissues (Bale, 1996; Danks, 2007).

Increasing cold-resistance can be triggered by environmental stimuli that also induce diapause (i.e. seasonal developmental arrest), such as the photoperiodic decrease, and in this case entering diapause also provides higher cold-resistance (Tauber et al., 1986; Koštál, 2006). In some species, increased cold resistance in winter is accomplished through an acclimation process (i.e. response to stress) when insects experience low temperatures (Lagerspetz, 2006). In such a case, acclimation is not necessarily associated with diapause (Hayward et al., 2014). The cause-effect relationship between winter diapause and cold-resistance in insects is still debated, and both phenomenon may not be directly linked in species or populations (Denlinger and Lee, 1991; Xie et al., 2015) (for a review on these links, see Hodkova and Hodek, 2004).

Diapause is generally expressed at one specific stage of the insect’s development (Tauber et al., 1986). Diapause during an immature stage has inherent metabolic costs because of the use of energetic reserves to withstand winter temperatures and survive throughout winter in an inactive stage (Schmidt and Conde, 2006; Hahn and Denlinger, 2011), and therefore involves a trade-off with some life-history traits at the adult stage (Kroon and Veenendaal, 1998; Fordyce et al., 2006). For instance, in the parasitoid *Asobara tabida*, the duration of larval diapause negatively impacts emerging females’ egg load after diapause (Ellers and Van Alphen, 2002). Larval diapause was also shown to be associated with a decrease in adult longevity in the parasitoid *Praon volucre* (Colinet et al., 2010).

Cold resistance itself, with or without diapause, can negatively affect different life-history traits such as longevity or heat-resistance in insects (Zera and Harshman, 2001; Hayward et al., 2005; Basson et al., 2012; Sulmon et al., 2015). Within insect populations, different thermal performance and overwintering strategies (diapause *vs.* non-diapause) can be observed, as a result of plastic responses to environmental variability such as bet-hedging (Hopper, 1999). Depending on the strategy used by an individual insect, trade-off in cold resistance could appear between life-stages when resources allocated at the diapausing immature stage become less abundant for the adult non-diapausing stage.

Aphidiinae aphid parasitoids (Hymenoptera: Braconidae) enter diapause as prepupae which allows measurement of potential trade-offs with the adults and to study costs and benefits of the diapause strategy. In populations from mild winter areas (e.g. Western Europe), diapause incidence has been decreasing for a few years and parasitoids of different species now mostly remain active throughout the year (Lumbierres et al., 2007; Gómez-Marco et al., 2015; Andrade et al., 2016; Tougeron et al., 2017). Exploring these trade-offs will help at understanding the rationale of changes in overwintering strategies in the context of climate warming. Parasitoids that do not enter diapause should thus be cold-resistant enough to withstand low temperatures during both their immature and adult stages. Parasitoids that remain active may have a fitness advantage because they avoid diapause associated costs while taking advantage at exploiting anholocyclic aphid populations present throughout the year. If so, selection should favor them over diapausing parasitoids, which could lead to a reduction in diapause expression in mild winter areas populations.

This study aimed at exploring the relationship between diapause and cold-resistance in a parasitoid population of *Aphidius ervi* from a mild winter area where both diapause and non-diapause strategies are expressed during winter (Andrade et al., 2016; Tougeron et al., 2017). We first hypothesized that energetic costs of diapause engender a trade-off between diapause at the prepupal stage and cold-resistance at the adult stage; adults that have been in prepupae diapause should be less cold-resistant than diapausing prepupae. We also expected adults that have not undergone diapause (i.e. adults that are active during winter) to have a higher fitness; they should be heavier and bigger than adults that come from diapausing prepupae. We then examined cold resistance in diapausing *vs.* non-diapausing individuals, as well as associated fitness consequences. We hypothesized that diapausing prepupae are more cold-resistant than non-diapausing ones, but due to trade-offs, adults that remain active in the winter should be more cold-resistant than adults that have undergone prepupal diapause.

## 2. Material & Methods

### 2.1. Biological material

A colony of the aphid *Sitobion avenae* F. (Hemiptera: Aphididae) was established from a single female collected at Le Rheu (France) in 1990. Aphids were reared on winter wheat sprouts (*Triticum aestivum*). Aphids were used as hosts for the parasitoid *A. ervi*, whose colony was established from individuals collected in 2015 around Rennes, France (48.1°N; 1.7°W). Both insects were reared at 20 ±1°C, 16:8 h Light:Dark (LD) photoregime and 70 ±10 % relative humidity (RH).

### 2.2. Diapause induction

Thirty < 48h-old, mated female parasitoids were split in two groups and introduced in two cages (50×50×50cm) with 350 to 400 *S. avenae* aphids of second or third instar for 48 h. Parasitoids were then removed from the cages and the cages with parasitized aphids were placed at 17°C, 10:14 h LD, 70% RH, conditions known to induce 8.3 ±4.2% diapause in our *A. ervi* population (Tougeron et al. 2017). Aphid mummification (i.e. dead aphids containing a parasitoid prepupa) was checked daily - starting 5 days after oviposition - and mummies were isolated in eppendorf tubes. This protocol was repeated three times over the course of the experiment (4 months) to obtain sufficient number of parasitoids to be tested.

Five different modalities were tested and produced as follow (Fig. 1); Less than 24 h after mummification, 27 mummies were taken to measure the SCP of the one-day-old prepupae (P1). Early parasitoid instars are the sensitive stage for diapause initiation in Aphidiinae (Brodeur and McNeil, 1989) but it was impossible to visually separate mummies containing diapausing or non-diapausing prepupae in the P1 treatment. We therefore tested the SCP of 21 one-day-old non-diapausing parasitoid prepupae formed at 20°C, 16:8h LD (P1C), a condition that does not induce diapause (Tougeron et al., 2017). Twenty-four adult parasitoids emerged up to 10 days after mummification were tested for their SCP within 12 h after their emergence (adults that have not undergone diapause, AND). Sixteen mummies from which no adults had emerged 15 days after mummification were considered to contain a 15-days-old diapausing prepupa (P15; Tougeron et al., 2017) and their SCP was measured. The remaining mummies were transferred at 20°C, 16:8 h LD to terminate diapause. The SCP of 25 adults emerging from this treatment was measured within 12 h after adult emergence (adults that went through diapause at their prepupae stage, AD). This latter treatment allowed us to measure the number of days required to break diapause in *A. ervi* under the experimental conditions tested (we stopped recording emergence 100 days after the mummies were placed at 20°C). To make sure parasitoids stayed in diapause long enough to see an effect on their SCP, we only tested individuals that emerged at least 40 days after being placed at 20°C.

**Figure 1:**
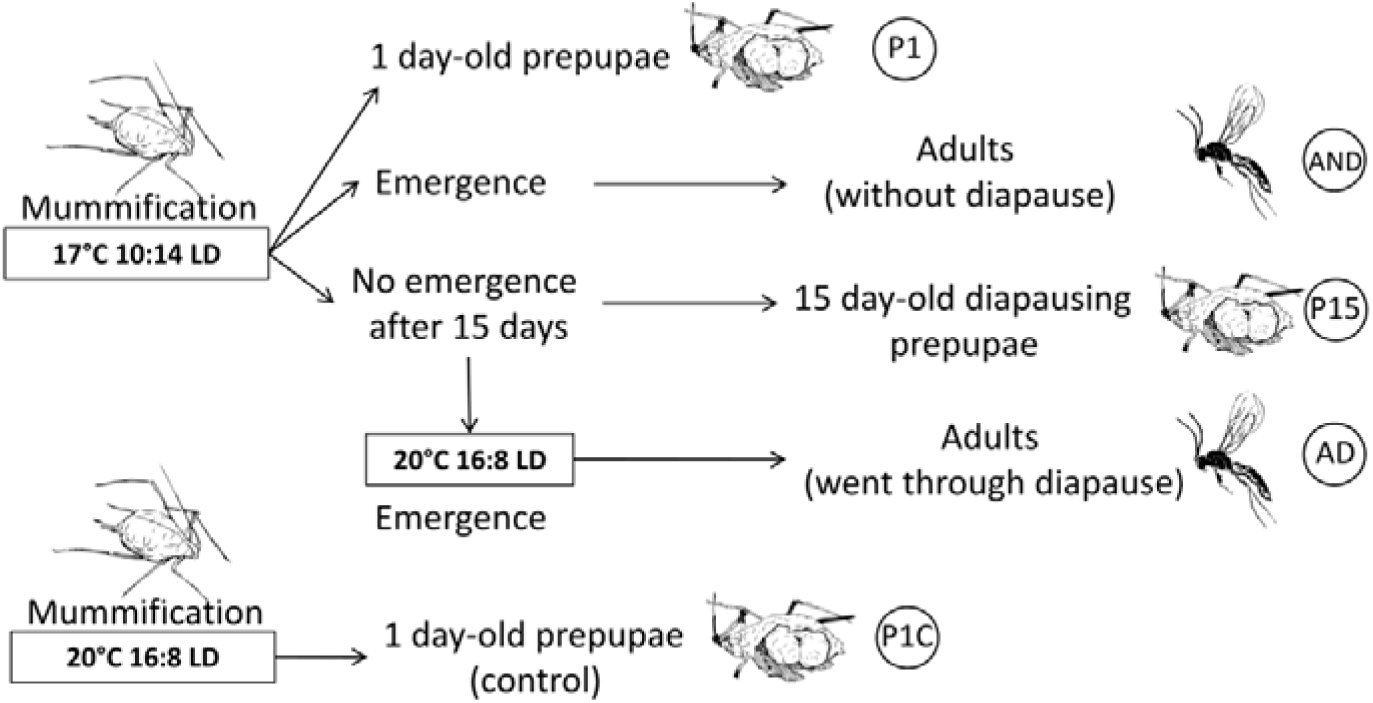
Schematic of the experimental design used to produce the five parasitoid treatments on which Super Cooling Points were measured. One-day-old prepupae in the mummy (P1), adults that did not enter prepupal diapause (AND), fifteen-days-old diapausing prepupae (P15), adults that went through prepupal diapause (AD), and one-day-old prepupae (Control; P1C).

### 2.3. SCP measurements

The use of SCP as a measure of cold resistance is controversial since mortality can occur long before the insect freezes. However, SCPs values are often well correlated to direct measures of cold resistance (e.g. LT50, CTmin), which makes SCP ecologically relevant to use (Cira et al., 2016; Renault et al., 2002), for instance in *Aphidius* parasitoids (Langer and Hance, 2000; Colinet et al., 2007).

Super Cooling Points of prepupae (within the mummy) and emerged adults were measured using metallic thermocouples plugged in a high resolution temperature data-logger (Testo 176-T4, ±0.3°C, Germany). Parasitoids were placed in a 7mm diameter gelatin capsule (Capsugel Coni-snap 1EL, NJ, USA) in direct contact with the thermocouple (Hanson and Venette, 2013). Capsules were then placed in glass test tubes on a rack that was immerged in the cryostatic bath. Temperature was set to cool at a rate of 0.2°C per minute, from +5°C to −30°C. Data was extracted from the data-logger using Testo Comfort Software 5.0. SCP corresponds to the onset of the exotherm produced by the latent heat of freezing, just before the death of the insect (Colinet et al., 2007; Renault et al., 2002).

### 2.4. Mass, volume and size measurements

We determined if the SCP was influenced by the mass of the parasitoids and the size of the adults or the volume of the mummies. Mass was measured using a ± 0.1 µg precision balance (Mettler-Toledo XP2U). Aphid mummies were weighted before and after each SCP experiment to test for potential effect of freezing on the mass. No difference was found (before SCP measurements: 0.36 ±0.04 mg, and after: 0.35 ±0.04 mg, Student paired t-test t=1.42, p=0.19, n=17), thus only the mass obtained after SCP measurements was kept for analyzes. Size of the adults and volume of the mummies were measured using a camera AxioCam ERc 5s (Zeiss, Germany) and the ImageJ software (v1.51). For adults, the length of the left hind tibia was measured. For mummies, length (L) and width (W) were measured and the volume (V) was obtained using the relation: *V = ΠLW*^2^/6 (Colinet et al., 2007). The mass/volume and mass/size ratio were then calculated and used in the analyses to determine if they had an influence on SCP. The sex of the adults was also determined as sexual dimorphism in size exists in *A. ervi* (Hurlbutt, 1987) and could influence the SCP (Renault et al., 2002).

### 2.5. Statistical analyses

A generalized linear model (GLM) was fitted to the data to test for differences in SCP among the five modalities (diapausing prepupae, non-diapausing prepupae, control prepupae, adult which went through immature diapause and adult which emerged without diapause) and using the sex and the ratio mass/size or mass/volume as covariables. Tukey post hoc tests for linear models (package ‘*multcomp*’) were then performed to test for differences among treatments. Data was then split between adults and prepupae and GLMs were fitted to the data in order to assess for differences in mass/size or mass/volume ratio between diapausing and non-diapausing parasitoids (sex was used as a covariable for adults). Differences between explanatory variables were assessed using a type-II Anova from the package ‘*car*’. All statistical analyses were performed using R software (R Core Team, 2017). The SCP distribution among the three prepupae treatment was plotted in order to make sure that no subgroup of prepupae (e.g. diapausing *vs.* non-diapausing) was observed within each treatment.

## 3. Results

Super Cooling Points differed significantly among parasitoid treatments (GLM, LR=75.6, df=4, p<0.001) (Fig. 2). The lowest SCP (i.e. the highest cold resistance) was found in the prepupae; both types of prepupae had similar SCPs (−24.2 ±0.3 °C and −24.7 ±0.4 °C for one-day, fifteen-days old diapausing prepupae, respectively, Tukey Contrasts, p=0.96). Their SCPs did not differ from the control group of one-day-old prepupae reared at 20 °C 16:8 h LD (−24.1 ±0.6 °C, Tukey Contrasts, p=0.97). The three types of prepupae had a lower SCP than adults that have not undergone diapause (−21.1 ±0.5 °C, Tukey Contrasts, p<0.05) and adults that have undergone diapause (−17.8 ±1.1 °C, Tukey Contrasts, p<0.05). Adults that have not undergone diapause had a higher cold-resistance than adults that went out of diapause (Tukey Contrasts, p<0.05).

**Fig 2:**
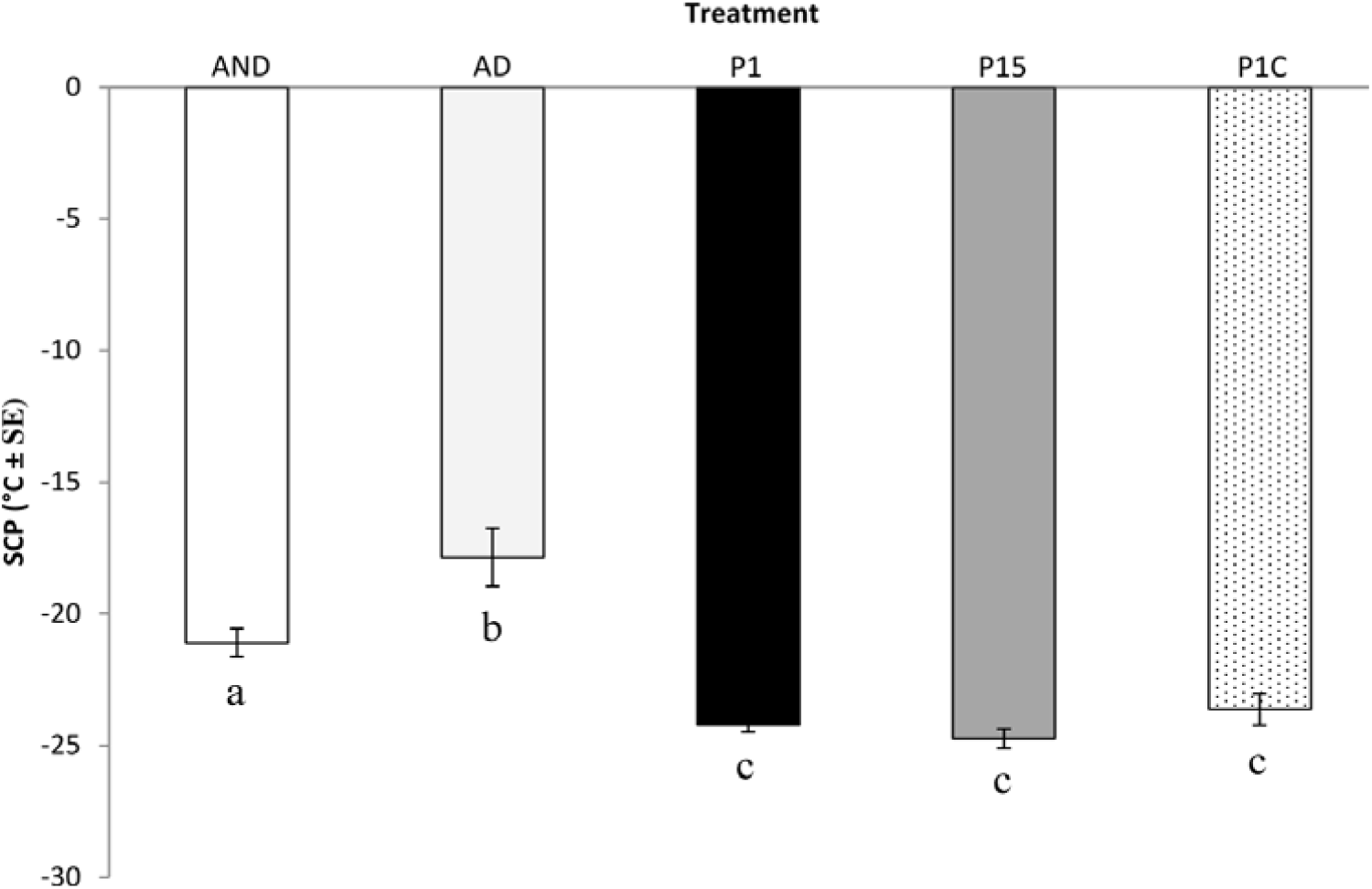
Differences in SCP (±SE) among parasitoid treatments; Adults that did not emerge from a diapausing prepupae (AND, n=24), adults that went through diapause at the immature instar (AD, n=25), one-day-old prepupae in the mummy (P1, n=27), fifteen-days-old diapausing prepupae in the mummy (P15, n=16), and control one-day-old prepupae in the mummy (P1C, n=21). Lowerscript letters indicate significant differences (p<0.05) between factors according to Tukey post-hoc tests for linear models following a GLM.

The distribution of SCP among prepupae treatments was homogeneous and unimodal. Even if both diapausing and non-diapausing prepupae could have been present in the P1 treatment, no SCP subgroup was observed.

From the prepupae placed at 20°C 16:8 h LD to break diapause, parasitoid adults took from 22 to 82 days to emerge (AD treatment) with a mean (±SE) emergence time of 61.4 ±3.3 days.

Mass/size ratios were smaller in AD (0.13 ±0.02) than in AND (0.22 ±0.02) and in P15 (0.26 ±0.06) than in P1 (0.47 ±0.06) (Fig. 3). This means that for a given body size, adults that have undergone diapause (AD) had a mass 1.6x lower than their non-diapausing counterparts (AND) (GLM, LR=7.0, df=1, p<0.01). Prepupae in diapause (P15) had a mass 1.8x lower than non-diapausing prepupae (P1) (GLM, LR=5.7, df=1, p<0.05) (Fig. 3).

**Fig 3:**
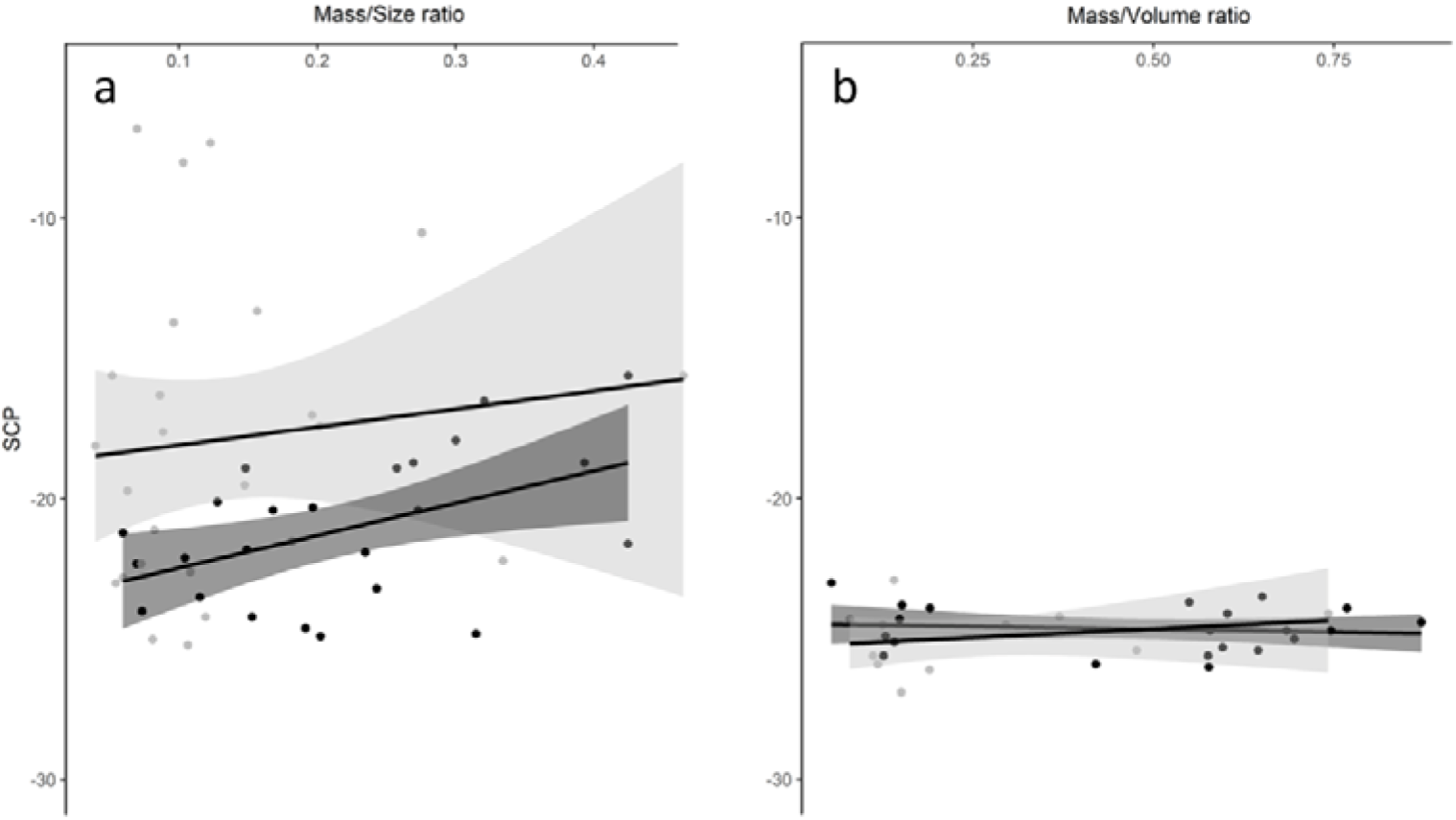
SCP (°C) variation depending on the (a) mass/size ratio of adults and (b) the mass/volume ratio of mummies, for diapausing prepupae (P15, n=16) or adults that come from diapause (AD, n=25) (grey) and non-diapausing prepupae (P1, n=27) or adults that did not undergo diapause (AND, n=24) (black). Lines and shaded areas show GLM predictions ± SE.

The mass/size ratio did not influence the SCP of adults (GLM, LR=2.6, df=1, p=0.10), for both AD and AND treatments (GLM interaction, LR=0.3, df=1, p=0.57), neither did the mass/volume ratio for prepupae (GLM, LR=0.005, df=1, p=0.98), for both P1 and P15 treatments (GLM interaction, LR=0.9, df=1, p=0.34) (Fig. 3). Sex had no effect on the SCP of the adults (GLM, LR=0.6, df=1, p=0.43).

There was SCP variability within treatments that could be explained by mass and size differences. When analyzed separately from the mass, small adults had a lower SCP than big ones (GLM, LR=6.2, df=1, p<0.05, R^2^=0.14) for both adults that have undergone diapause and adults that have not undergone diapause (interaction, GLM, LR=0.03, df=1, p=0.84). Moreover, when analyzed separately from the size, light adults had a lower SCP than heavy ones (GLM, LR=4.4, df=1, p<0.05, R^2^=0.05), for both adult treatments (interaction, GLM, LR=0.58, df=1, p=0.44). This relationship was not true for the volume (GLM, LR=1.8, df=1, p=0.18) and the mass (GLM, LR=0.96, df=1, p=0.33) of the prepupa treatments.

## 4. Discussion

*Aphidius ervi* adults that emerged from individuals that went through diapause had a lower cold resistance (i.e. a higher SCP) than any parasitoids from the other treatments suggesting a trade-off in cold-resistance within and between life-stages, depending on the diapause *vs.* non-diapause strategy used by the parasitoid. Our results also suggest that the non-diapause strategy could be advantageous in mild winter areas because parasitoids are able to withstand winter temperatures while avoiding diapause costs. Unexpectedly, diapausing prepupae did not have a higher cold resistance than non-diapausing prepupae, since no SCP differences were observed between 1-day-old and 15-days-old (diapausing) prepupae.

Our results suggest a trade-off between immature (prepupa) and adult cold-resistance when parasitoids enter diapause. This phenomenon might be the result of energetic expenses during diapause, leading to less energetic reserves to sustain low temperatures for the emerging individual. We also found that adult parasitoids that have undergone diapause and diapausing prepupae were lighter than their non-diapausing counterparts, highlighting a physiological cost to diapause. It is known in some parasitoid species that the fitness of adults emerging following a diapause episode decreases with diapause duration (Ellers and Van Alphen, 2002; Ito, 2007). We also highlighted costs in term of physiological functions as shown by the lower SCP of adults that went through larval diapause. The observed trade-off in cold-resistance is consistent to the fact that parasitoids terminate diapause when favorable conditions return. In this case, energy could be allocated to sustain other functions than thermal resistance.

*Aphidius ervi* exhibited SCP values in the range of values observed in closely related species (Langer and Hance, 2000; Colinet et al., 2007) but no SCP difference was found between the three types of prepupae in the present study. In aphid parasitoids, the relation between SCP and diapause is not yet clear, as the SCP of immature instars of *A. ervi* and *Aphidius rhopalosiphi* was found to be low during diapause (i.e. high cold-resistance) (Langer and Hance, 2000) while no link was found in *Praon volucre* (Colinet et al., 2010). First, excretion of nucleating gut content (i.e. meconium) after the nymphal metamorphosis may have limited action of cold on both types of prepupae in a similar fashion. Second, cold-hardening may happen later in the process of diapause (after 15 days), as parasitoids can remain in diapause over several months in natural conditions (Koštál, 2006). In addition, higher cold-resistance may appear only if insects are exposed to cold temperatures, following acclimation in winter, either during diapause or not (Denlinger and Lee, 1991; Hodkova and Hodek, 2004; Lester and Irwin, 2012). In our study, parasitoids from each treatment developed and acclimated at the same temperature.

Finally, in insects, the relation between supercooling temperature and sex or morphological traits such as size or mass is still in debate. Colinet et al. (2007) showed that the more important the volume of the mummy, the less cold resistant *Aphidius colemani* prepupae in the mummy was. The ability to supercool (i.e. to resist crystallization) decreases as mass, size and volume increase (Colinet et al., 2007). Such a pattern was not observed in our *A. ervi* population. This may be due to interspecific differences or to different rearing photoperiods (16:8 h LD in Colinet et al. (2007) and 10:14 h LD in our study) as it was shown to influence morphological traits, development times and insects’ fitness independently of the temperature (Joschinski et al., 2015).

Sexual dimorphism in size, mass and energetic reserves have to be carefully considered when exploring differences in thermal tolerance (Blanckenhorn, 2000; Le Lann et al., 2011; Ismail et al., 2012). In adults, when taken apart from each other, size and mass influenced the SCP of the parasitoids. This is in line with previous studies on parasitoids showing that small individuals are more cold-resistant than large parasitoids (Ismail et al., 2012; Tougeron et al., 2016), due to more parsimonious use of energetic reserves (Reim et al., 2006). Differential resistance to cold between sexes has been reported in aphid parasitoids (Le Lann et al., 2011), with adult females being less resistant to cold than males, probably due to size differences between sexes. We observed no such difference in our study, suggesting that Aphidiinae parasitoids’ SCP is influenced by the size and not by sex *per se*.

The population of *A. ervi* used in this study does not express high levels of diapause (mean 11.2 ± 4.9% at 14 °C, 10:14 h LD; Tougeron et al., 2017), and parasitoids mostly remain active as non-diapausing individuals throughout winter (Andrade et al., 2016), suggesting a risk-spreading strategy. The relative proportions of diapausing and non-diapausing parasitoids is likely to be determined by the risk of encountering cold spells or harsh winters at a given location (Hopper, 1999). Parasitoids can either (i) overwinter as diapausing prepupae, and then resist winter cold but miss opportunities to exploit a favorable winter environment and have a lower fitness as emerging adults, or (ii) abort diapause and be better cold-resistant at the adult stage to withstand possible cold spells and exploit the winter environment. This latter strategy may be advantageous and thus selected if diapausing prepupae does not have a better cold resistance than non-diapausing ones, and if parasitoids are cold-resistant enough to overwinter as adults, which seems to be the case in mild-winter areas such as western Europe (Lumbierres et al., 2007; Andrade et al., 2016; Tougeron et al., 2016, 2017).

## 5. Conclusion

To conclude, diapause expression could be reduced if the balance between its physiological (e.g. energetic demand) or ecological costs (e.g. no exploitation of the winter environment, less generation produced each year) and its benefits (e.g. increased cold resistance, synchronization with the host) is shifted by climate warming (Bale and Hayward, 2010; Sgrò et al., 2016). In more natural setting, insects can adopt behavioural adjustments of their thermal tolerance capacities, which may participate in adjusting their response to climate warming (Sunday et al., 2014). For instance, parasitoids are known to manipulate their host in the context of diapause to make them choose sheltered areas in which they can overwinter (Brodeur and McNeil, 1989b). Non-diapausing parasitoids and their hosts can also live in microclimatic habitats (thermal shelters) that can modify their thermal tolerance capacities (Tougeron et al., 2016; Alford et al., 2017). There is now a need to better understand the winter ecology and physiology of non-diapausing insects, as they are facing increasing winter temperatures, especially in the context of climate warming (Owen et al., 2013; Stuhldreher et al., 2014).

## Acknowledgments

We thank S. Llopis for technical assistance and N. Bidet for drawing insects in Fig. 1. KT was supported by the French Région Bretagne and the Canada Research Chair in Biological Control. LB and KT performed the experiments and analyzed the results. KT wrote the manuscript. All co-authors contributed to the protocols and made substantial revisions to the manuscript. This study was supported by the LTER France Zone Atelier Armorique.

